# Epithelial Immunomodulation by Aerosolized Toll-like Receptor Agonists Attenuates Allergic Responsiveness in Mice

**DOI:** 10.1101/2021.08.05.455337

**Authors:** David L. Goldblatt, Gabriella Valverde Ha, Shradha Wali, Vikram V. Kulkarni, Michael K. Longmire, Ana M. Jaramillo, Rosha P. Chittuluru, Adrienne Fouts, Margarita Martinez-Moczygemba, Jonathan T. Lei, David P. Huston, Michael J. Tuvim, Burton F. Dickey, Scott E. Evans

**Affiliations:** Department of Pulmonary Medicine, University of Texas MD Anderson Cancer Center, Houston, Texas, 77030, USA; Department of Microbial and Molecular Pathogenesis, Texas A&M Health Science Center, Houston, Texas, 77030, USA; Clinical Science and Translational Research Institute, Texas A&M Health Science Center, Houston, Texas, 77030, USA; Howard Hughes Medical Institute, Chevy Chase, MD, 20815 USA; University of Texas Rio Grande Valley School of Medicine, Edinburg, TX, 78539

## Abstract

Allergic asthma is a chronic inflammatory respiratory disease associated with eosinophilic infiltration, increased mucus production, airway hyperresponsiveness (AHR), and airway remodeling. Epidemiologic data has revealed that the prevalence of allergic sensitization and associated diseases has increased in the twentieth century. This has been hypothesized to be partly due to reduced contact with microbial organisms (the hygiene hypothesis) in industrialized society. Airway epithelial cells, once considered a static physical barrier between the body and the external world, are now widely recognized as immunologically active cells that can initiate, maintain, and restrain inflammatory responses, such as those that mediate allergic disease. Airway epithelial cells can sense allergens via myriad expression of Toll-like receptors (TLRs) and other pattern-recognition receptors (PRRs). We sought to determine whether the innate immune response stimulated by a combination of Pam2CSK4 (“Pam2”, TLR2/6 ligand) and a class C oligodeoxynucleotide ODN362 (“ODN”, TLR9 ligand) when delivered together by aerosol (“Pam2ODN”), can modulate the allergic immune response to allergens. Treatment with Pam2ODN 7 days before sensitization to House Dust Mite (HDM) extract resulted in a strong reduction in eosinophilic and lymphocytic inflammation. This Pam2ODN immunomodulatory effect was also seen using Ovalbumin (OVA) and A. oryzae (Ao) mouse models. The immunomodulatory effect was observed as much as 30 days before sensitization to HDM, but ineffective just 2 days after sensitization, suggesting that Pam2ODN immunomodulation lowers the allergic responsiveness of airway epithelial cells. Furthermore, Pam2 and ODN cooperated synergistically suggesting that this treatment is superior to any single agonist in the setting of allergen immunotherapy.

**One Sentence Summary:** A synergistic combination of Toll-like Receptor agonists, delivered directly into the lung mucosa, can attenuate allergic responsiveness of airway epithelial cells and prevent host sensitization to aeroallergens.

**What is already known:**

- Allergic sensitization has increased in the 20^th^ century due to reduced contact with microbial organisms in industrialized society (ie. hygiene hypothesis)
- We have previously identified a pharmacological means to stimulate innate immunity of lung epithelial cells.

**What this study adds:**

- Activation of innate immunity in lung epithelial cells attenuates the allergic responsiveness of mice.
- Synergistic cooperation of pattern recognition receptors induces stronger immunomodulatory responses

**What is the clinical significance:**

- Aerosolized Toll-like Receptor agonists have been demonstrated as safe in human clinical trials
- This study provides proof-of-principle that aerosolized toll-like receptor agonists could have clinical efficacy in the setting of the allergen immunotherapy

## Introduction

Allergic asthma is a chronic inflammatory respiratory disease associated with eosinophilic infiltration, increased mucus production, airway hyperresponsiveness (AHR), and airway remodeling.^1^ Epidemiologic data has revealed that the prevalence of allergic sensitization and associated diseases has increased in the twentieth century. This has been hypothesized to be partly due to reduced contact with microbial organisms (the hygiene hypothesis) in industrialized society^2^. In one study comparing Amish and Hutterite children, higher levels of microbial elements in farm dust were found to be strongly protective against developing asthma.^3^ In mechanistic studies, this effect was found to be mediated by downregulation of inflammatory pathways within airway epithelial cells.^4^

Airway epithelial cells, once considered a static physical barrier between the body and the external world, are now widely recognized as immunologically active cells that can initiate, maintain, and restrain inflammatory responses, such as those that mediate allergic disease.^5^ Lung airway epithelial cells express myriad pattern recognition receptors (PRRs) such as Toll-like receptors (TLRs) that enable them to sense and respond to a variety of external triggers, usually referred to as pathogen-associated molecular patterns (PAMPs). The epithelial-derived cytokines IL-33, thymic stromal lymphopoietin (TSLP), and IL-25 are widely recognized as alarmins produced in response to allergens, such as those associated with house-dust mites (HDM). These cytokines activate leukocytes involved in allergic inflammation, such as eosinophils, T helper 2 (T_H_2) cells, mast cells, basophils, type 2 innate lymphoid cells (ILC2), and dendritic cells (DC), which work together to polarize the immune response in a type 2 direction.

Our group has studied how to stimulate lung epithelial cells to defend against microbial pathogens.^6^ We have shown that a combination of Pam2CSK4 (“Pam2”, TLR2/6 ligand) and a class C oligodeoxynucleotide ODN362 (“ODN”, TLR9 ligand), when delivered together by aerosol (“Pam2ODN”) synergistically activate an innate immune response in the lung mucosa (but not systemically) which results in host resistance to bacteria, fungi, and viruses.^7–10^ Pam2ODN-mediated pathogen resistance is initiated very rapidly and has been shown to be mediated by lung epithelial cells.^11^

More recently, we showed that Pam2ODN can attenuate chronic asthma-like lung disease in mice infected with Sendai virus (SeV).^12^ We reasoned that Pam2ODN would exert a protective effect against the chronic disease by reducing the viral burden in the acute infection, but we observed some efficacy when Pam2ODN treatment was given at time points remote from the viral challenge and did not reduce the virus burden. This partial discordance between the reduction in viral burden and chronic disease severity led us to hypothesize that Pam2ODN also modulated the type 2 immune response induced by the virus infection that drives the chronic inflammatory disease.

To disentangle antimicrobial and immunomodulatory effects of Pam2ODN, we now use allergic models where the antimicrobial effect is irrelevant. In this study, we show that Pam2ODN attenuates the type 2 allergic immune response to HDM, Ovalbumin (OVA), and *Aspergillus oryzae* (Ao) in the lung mucosa.

## Results

### Pam2ODN prevents allergic inflammation to HDM

Mice were sensitized by airway instillation of 100 μg HDM extract at day 0, followed by 6 daily challenges of 10 μg HDM extract from day 7 to 12, and evaluation of allergic inflammation on day 15 (Fig. 1A).^13^ As previously reported, HDM-sensitized, HDM-challenged (HDM/HDM) mice exhibited robust allergic inflammation, though PBS-sensitized, HDM-challenged (PBS/HDM) did not (Fig. S1). Allergic inflammation to HDM was reflected by the substantial influx of eosinophils (Fig 1B, orange arrowhead) and lymphocytes (Fig. 1B, black arrowhead) with small numbers of neutrophils (Fig 1B, blue arrowhead). Macrophages were increased only slightly in number but were observed to be larger and more intensely stained than PBS/PBS, indicating activation (Fig. 1B, open arrowhead).

**Figure 1.**
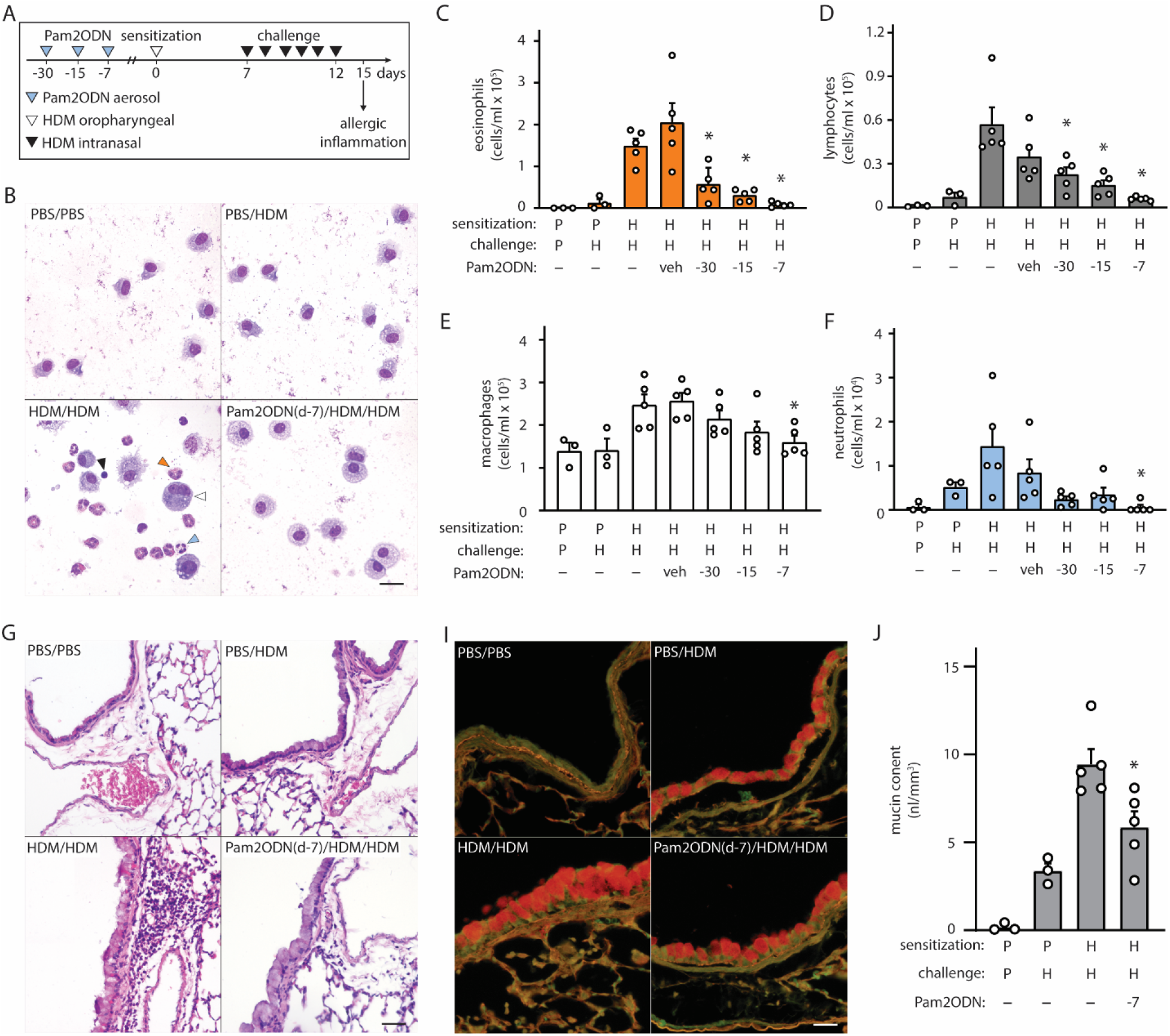
Exposure to Pam2-ODN attenuates allergic inflammation to HDM. **(A)** HDM experimental paradigm with a sensitization of 100 µg HDM and challenges of 10 µg HDM. **(B)** Leukocytes obtained by lung lavage were pelleted onto glass slides by centrifugation and stained with Wright-Giemsa. Scale bar = 20 μm. **(C-F)** Quantification of leukocytes in lung lavage for **(C)** eosinophils, **(D)** lymphocytes, **(E)** macrophages, and **(F)** neutrophils (N = 3-5 mice). **(G-H)** Airways stained with H&E to demonstrate submucosal inflammation. Scale bar = 40 μm **(I)** Airway epithelium stained with PAFS to demonstrate intracellular mucin in red. Scale bar = 20 μm. **(J)** Quantification of intracellular mucin content by image analysis of airway (N = 3-5 mice). Bars show mean +/- SEM. P* < 0.05 by one-way ANOVA with Dunnett’s test for multiple comparisons against [HDM/HDM] control. P = PBS. H = HDM.

When mice were treated with a single Pam2ODN treatment 7 days before HDM sensitization, we observed large reductions in leukocytes collected in lung lavage fluid, compared to HDM/HDM mice (Fig. 1B). Differential cell quantification with Wright-Giemsa stain revealed a >90% decrease in eosinophils (Fig. 1C), >85% decrease in lymphocytes (Fig. 1D), > 30% decrease in macrophages (Fig 1E), and > 95% decrease in neutrophils (Fig. 1F). Pam2ODN treatments similarly showed efficacy when administered 30- and 15-days before sensitization, with a trend toward less efficacy when the time between treatment and sensitization was increased.

On H&E staining of lung tissue, most of the infiltrating leukocytes were localized to the submucosal space between major airways and vasculature, though some alveolar inflammation was also observed (Fig. 1G-H, Fig. S2). Mucous metaplasia of lung epithelial cells was assessed by staining lung tissue with fluorescent PAS stain (PAFS) (Fig. 1I).^14^ Quantitative image analysis of intracellular mucin content revealed that intracellular mucin accumulates moderately in PBS/HDM mice despite the near-complete absence of inflammatory leukocytes, though HDM sensitization was required for the full phenotype (Fig. 1J). A single Pam2ODN treatment 7 days before HDM sensitization reduced intracellular epithelial mucin content >35%.

### Pam2ODN blocks sensitization to HDM in the airway

We next evaluated whether Pam2ODN prevented sensitization to HDM or attenuated allergic inflammation during the challenge phase. To do this, we varied the time-interval and the relative chronological order of Pam2ODN treatment and HDM sensitization. When Pam2ODN treatment was administered just 1 day before HDM sensitization, eosinophils were still significantly reduced, but this effect was completely absent when treatment was administered just 2 days after HDM sensitization, and at any later time point (Fig 2A). Since multiple treatments of Pam2ODN had higher efficacy than a single treatment (Fig. S3), we evaluated whether the lack of Pam2ODN efficacy after HDM sensitization could be due to insufficient treatment by administering 6 daily Pam2ODN treatments after HDM sensitization. As was observed using single treatments, multiple treatments after HDM sensitization did not reduce allergic inflammation (Fig. 2B).

**Figure 2.**
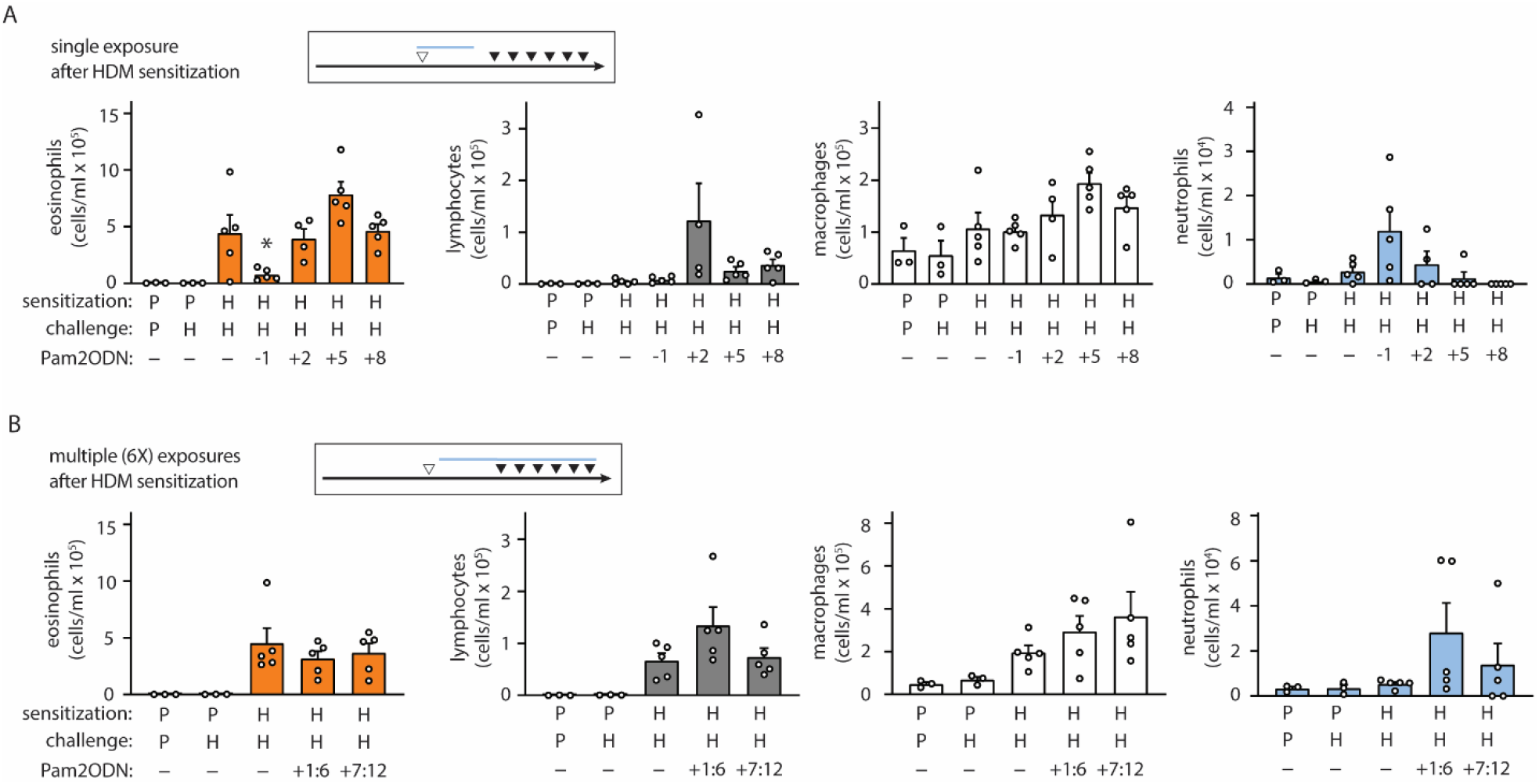
Pam2-ODN prevents sensitization to HDM. **(A-B)** Quantification of leukocytes in lung lavage when a single Pam2-ODN treatment was administered before or after sensitization to HDM (N = 3-5 mice). **(B)** Quantification of lung leukocytes when 6 consecutive daily Pam2-ODN treatments were administered after sensitization to HDM during the first week (days +1:6) or concurrently with challenge (days +7:12). Pictograms show blue bar at the approximate range of Pam2-ODN exposure in relationship to sensitization and challenge. Bars show mean +/- SEM. P* < 0.05 by one-way ANOVA with Dunnett’s test for multiple comparisons against [HDM/HDM] control. P = PBS. H = HDM.

We also evaluated whether Pam2ODN could prevent allergic inflammation when HDM sensitization occurred at a different location than the lungs. To do this, we used the same time schedule for HDM sensitization and challenge as previously shown, except that HDM sensitization was performed by intraperitoneal injection of 100 μg HDM (Fig S4A). Pam2ODN treatments were administered at day 0 and day 6 to cover the entire challenge period, however this resulted in no significant reduction in allergic inflammation (Fig. S4B-E).

### Pam2 and ODN interact synergistically

The ligands Pam2 and ODN were selected for their remarkable synergy, however this characteristic has only been evaluated for antimicrobial resistance thus far.^7^ To evaluate whether Pam2 and ODN also cooperate for allergic immunomodulation, the agonists were aerosolized individually or in combination 7 days before HDM sensitization. Neither Pam2 nor ODN exhibited any significant efficacy when administered alone, but the combination Pam2ODN treatment completely blocked allergic inflammation (Fig 3A-D).

**Figure 3.**
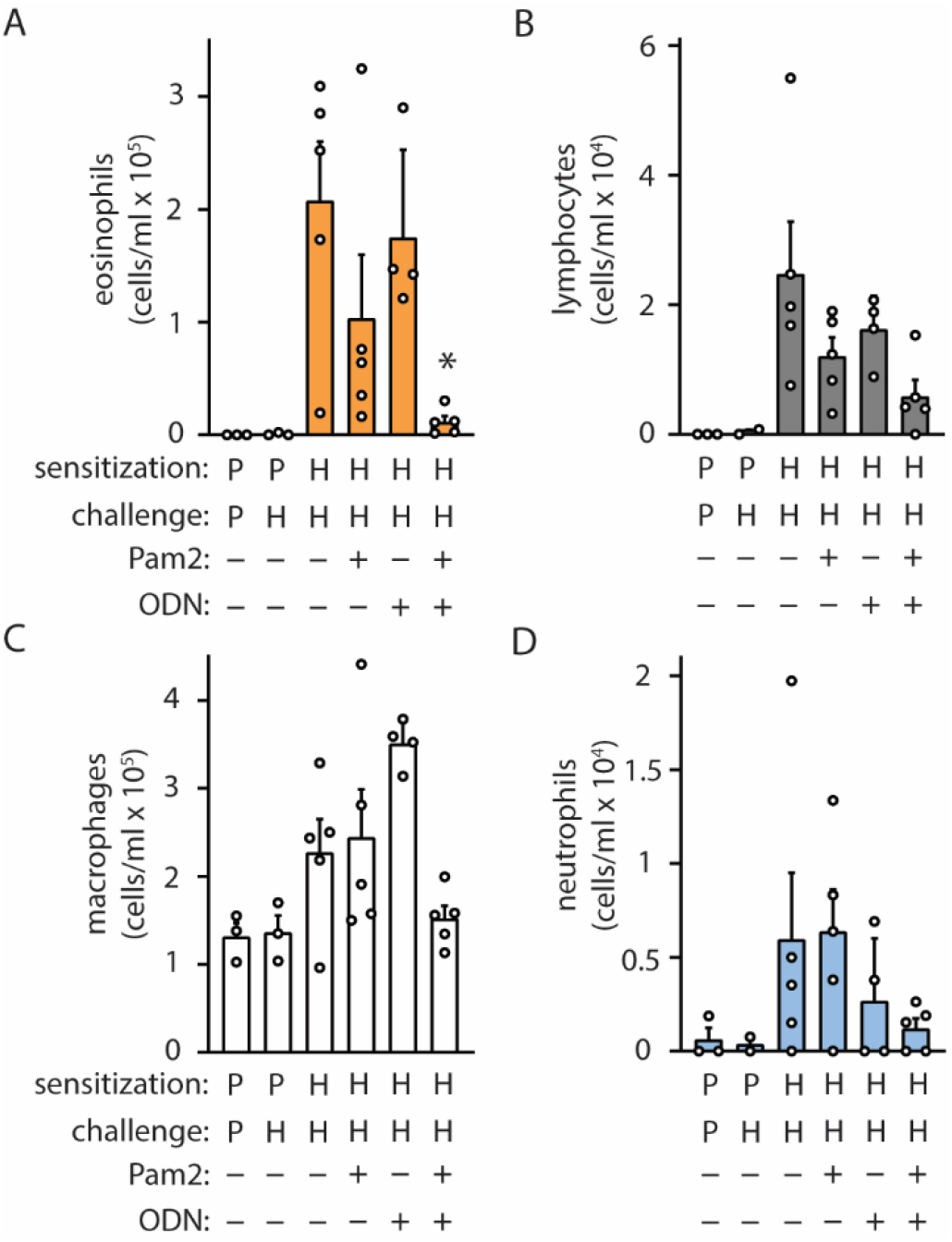
Pam2 and ODN interact synergistically to block sensitization to HDM. Pam2, ODN, or Pam2ODN was administered 7 days before sensitization to HDM. Quantification of leukocytes in lung lavage for **(A)** eosinophils, **(B)** lymphocytes, **(C)** macrophages, and **(D)** neutrophils (N = 3-5 mice). Bars show mean +/- SEM. P* < 0.005 for synergy between Pam2 and ODN by linear regression. P = PBS. H = HDM.

### Pam2ODN attenuates allergic inflammation to OVA

To evaluate the generalizability of Pam2ODN allergic immunomodulation, we also tested a conventional 3-week OVA model (Fig. 4A), consisting of 2 sensitizing intraperitoneal injections during the first week, and aerosol challenges beginning on day 8 administered every 2-3 days until day 21. From day 21 to day 24, OVA aerosol challenges were given daily, and allergic inflammation was assessed at day 24. Pam2ODN treatments were given immediately before OVA challenges. OVA-sensitized, OVA-challenged (OVA/OVA) mice had elevated cell counts in eosinophils, lymphocytes, and neutrophils, compared to PBS-sensitized, OVA-challenged (PBS/OVA) mice (Fig 4B-E). Pam2ODN treatment reduced eosinophils >75%, but macrophages and lymphocytes were unchanged. With this Pam2ODN treatment model, neutrophils were observed to be elevated, however this is likely a direct response to Pam2ODN, which elicits neutrophilic inflammation over a period of 72 hours, rather than a secondary effect on the allergic response.^15^ Airway epithelial mucin content was also decreased >25% by Pam2ODN treatment (Fig. 4F).

**Figure 4.**
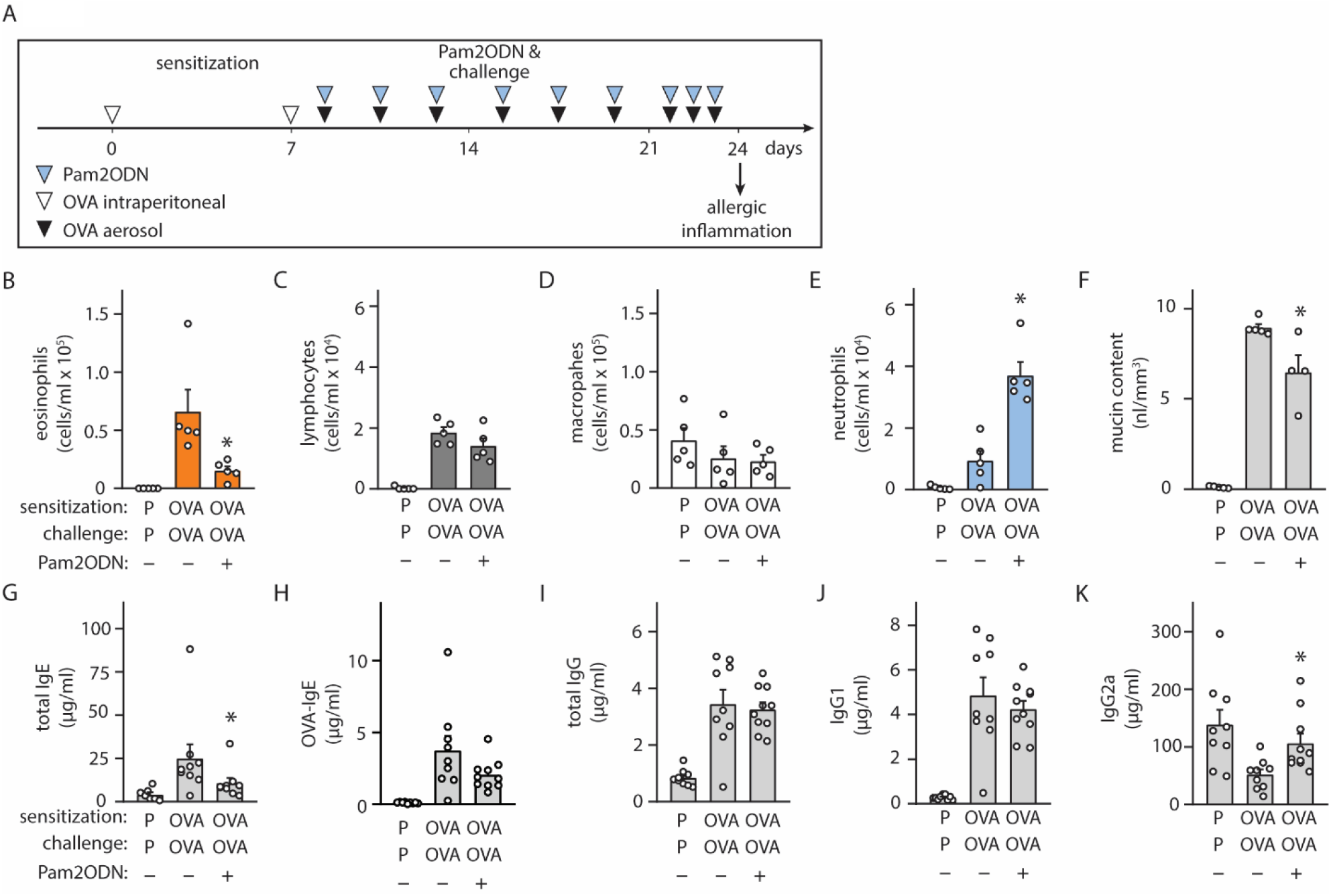
Exposure to Pam2-ODN attenuates allergic inflammation to OVA. **(A)** OVA experimental paradigm with sensitization of 20 µg OVA by intraperitoneal injection and aerosol challenges of 2.5% OVA. **(B-E)** Quantification of leukocytes in lung lavage for **(B)** eosinophils, **(C)** lymphocytes, **(D)** macrophages, and **(E)** neutrophils (N = 4-5 mice). **(F)** Quantification of intracellular mucin content by image analysis of airway stained with PAFS (N = 4-5 mice). **(G-K)** Quantification of serum immunoglobulin concentrations by ELISA for **(G)** IgE, **(H)** OVA-specific IgE, **(I)** IgG, **(J)** IgG1, and **(K)** IgG2a (N = 8-9 mice). Bars show mean +/- SEM. P* < 0.05 by unpaired students’ T test for OVA vs OP-OVA. P = PBS. OVA = Ovalbumin.

Assessing changes in the systemic immune response, Pam2ODN treatment reduced serum immunoglobulin E (IgE) levels by >50% (Fig. 4G), including a strong trend towards reducing OVA-specific IgE levels (Fig. 4H), but total IgG (Fig. 4I) and IgG1 (Fig. 4J) were unchanged. IgG2a levels, which increase with type 1 immune responses, decreased in OVA/OVA mice, compared to PBS/PBS, and Pam2ODN treatment reversed this change (Fig. 4K).

### Pam2ODN attenuates allergic inflammation to Aspergillus oryzae

Allergic sensitization can occur following exposure to fungal antigens.^16^ To evaluate whether fungal allergic inflammation could also be attenuated by Pam2ODN, we tested a combined Ao-OVA (AoO) model (Fig. 5A), with a similar time schedule as with OVA alone (Fig. 4A), except Ao-OVA or Pam2ODN were not administered from days 21 to 24. Pam2ODN treatment caused a >50% reduction in eosinophils (Fig 5B). Macrophages were decreased in AoO-challenged mice, and this was reversed by Pam2ODN treatment (Fig. 5C). Lymphocytes were not significantly elevated in AoO challenged mice, compared to mice only challenged with PBS (Fig. 5D). Neutrophil levels were not increased in Pam2ODN-treated groups (Fig 5E) due to the removal of Pam2ODN treatments within 3 days of evaluation (Fig. 5A). Airway epithelial mucin content was also observed to be decreased >30% with Pam2ODN treatment (Fig. 5F).

**Figure 5.**
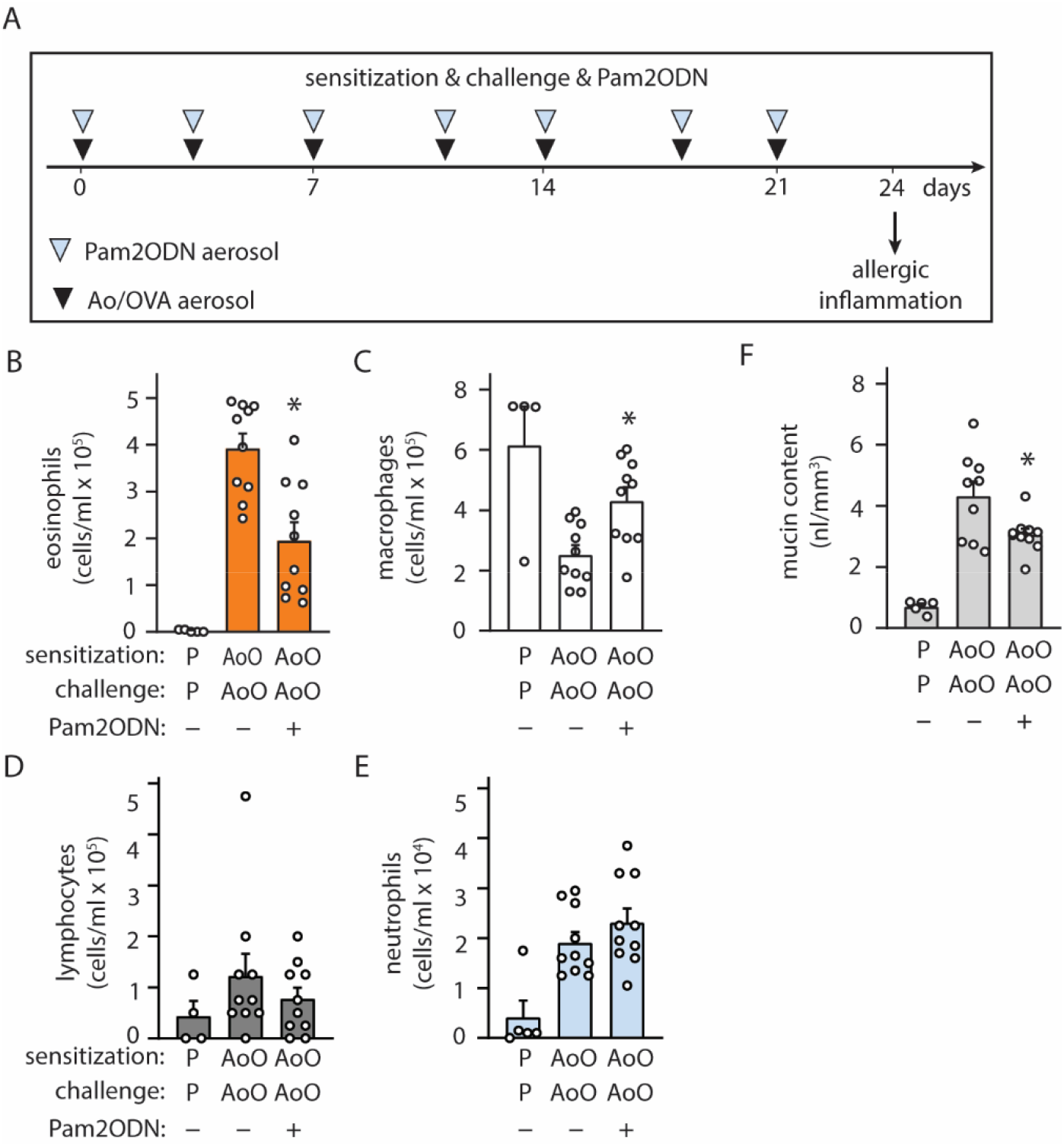
Exposure to Pam2-ODN prevents allergic inflammation to Aspergillus oryzae. **(A)** Ao-OVA experimental paradigm with aerosol challenges of 10 µg Ao and 2.5% OVA. **(B-E)** Quantification of leukocytes in lung lavage for **(B)** eosinophils, **(C)** lymphocytes, **(D)** macrophages, and **(E)** neutrophils (N = 4-9 mice). **(F)** Quantification of intracellular mucin content by image analysis of airway stained with PAFS (N = 4-9 mice). Bars show mean +/- SEM. P* < 0.05 by unpaired students’ T test for OVA vs OP-OVA.

## Discussion

Over the last two decades, scientific interest in utilizing PRR agonists, particularly TLRs, for the development of novel rapid-onset immunotherapy has grown considerably.^17^ However, these novel agents have not yet achieved significant clinical success, as exemplified by a recent study using a TLR9 agonist that failed to improve asthma control.^18^ The inability to translate preclinical success into clinical results is likely due to the complexity of allergic sensitization and the interplay of different variables affecting the degree of type 2 polarization. Our study has identified a novel combination of TLR agonists that cooperate synergistically to modulate the immune response to aeroallergens, offering the potential for more robust immunomodulatory effects than those previously tested.

In an earlier study, we found that Pam2ODN had immunomodulatory effects on the chronic asthma-like disease that persists after SeV viral clearance that could not be explained by Pam2ODN-mediated antiviral properties.^12^ In the current study, we observed Pam2ODN was effective at reducing allergic inflammation and mucous metaplasia induced by three different allergens: HDM, OVA, and Ao. While each model displays unique immunogenic characteristics, SeV, HDM, OVA, and Ao all lead to a chronic lung disease driven by polarization of the lung mucosa towards type 2 immunity. The effectiveness of this treatment against all antigens evaluated, as well as the immunomodulatory effects on the SeV-induced chronic disease from our previous study,^12^ suggests that Pam2ODN is a broadly effective immunomodulatory agent that can regulate the type 2 immune response.

One of the most striking results is the speed of onset for Pam2ODN-induced immunomodulation, with onset of efficacy observed in as little as 24 hours before HDM sensitization. Leukocyte recruitment to the lung by Pam2ODN has been measured previously to reach a peak 48 hours after exposure to Pam2ODN,^15^ and thus is unlikely to be responsible for the immunomodulatory effect seen at this early time point. On the other hand, epithelial cytokine release following Pam2ODN exposure has been observed within 4 hours.^15^ Additionally, it has been well established that HDM sensing by lung epithelial cells, primarily via TLR4, is required for the development of the full allergic phenotype.^19^ We further show that lung epithelial cells can respond to HDM directly and immediately by the observation that PBS/HDM mice show accumulated intracellular mucin and no detectable eosinophil recruitment. This indicates that lung epithelial cells can generate an allergic response directly and that this can occur before the innate or adaptive leukocyte-mediated immune responses have developed. Taken together, these data suggest that integration between the type 2 immunogens and Pam2ODN immunomodulation takes place within lung epithelial cells, which are optimally positioned to be first responders to both inputs.

By varying single treatments of Pam2ODN at different time points before and after HDM sensitization, we were able to show that Pam2ODN treatment is only effective when administered before HDM sensitization. We observed Pam2ODN efficacy in as little as 24 hours before sensitization, but a single treatment 2 days after sensitization was completely ineffective. These results suggest that once the host immune response to allergens has begun to polarize in a type 2 direction it is very difficult to modulate in another direction.

Considering the results from the HDM model in isolation, it would be difficult to determine whether Pam2ODN prevented host sensitization by modulating airway responses or systemic pathways downstream. However, since Pam2ODN is delivered directly to the airway mucosa via aerosol and exerts virtually no effect on systemic immune responses,^15^ this suggests that primary action on systemic immune responses would be highly implausible. As we also observed efficacy in our intraperitoneally sensitized OVA model, this supports our inference that Pam2ODN treatment modulates the allergic responsiveness of the airway mucosa and provides a rationale for treatment in already systemically sensitized hosts.

Interestingly, we also observed an overall change toward less systemic immunity to OVA in Pam2ODN-treated mice, reflected by lower levels of total IgE, OVA-specific IgE and IgG2a. This suggests that airway modulation during repeat exposures to allergen may block systemic re-sensitization, despite the presence of allergen-specific T cells. This could mean that a long treatment protocol with Pam2ODN might prevent the progression of atopic sensitivity.

We also identified that the immunogenicity of systemically sensitized leukocytes can vary significantly, and if sufficiently strong, can overcome the Pam2ODN-induced immunomodulation. We developed a novel adaptation of the acute HDM model that used intraperitoneal injection of HDM extract, followed by airway HDM challenges. within contrast to our observations in the OVA model, Pam2ODN airway immunomodulation was ineffective at preventing airway allergic polarization. One possible explanation for this discord is that HDM is an intrinsically stronger type 2 immunogen than OVA, and that the HDM-specific T cells provide a stronger pro-Th2 stimulus than OVA-specific T cells, and that Pam2ODN immunomodulation is not equally as strong to provide a counter-regulatory stimulus. HDM immunogenicity strength is further supported by the previous observations that HDM extract is capable of sensitizing hosts directly in the lung mucosa and does not require pairing with an adjuvant. This hypothesis can be explicitly tested in future work.

Because different allergens appear to have different immunogenic strength, it is plausible that effective immunomodulation of strong immunogens require equivalently strong immunomodulatory stimulus. Any single PRR agonist will ultimately be restricted by receptor saturation, and thus cannot invoke a stronger response above a set threshold concentration. However, Pam2ODN provides an important clue to bypassing this immunomodulatory strength limit by pairing immunomodulatory agents that exhibit synergy. Previous work has shown that Pam2 and ODN cooperate synergistically to mediate protection against microbial pathogens, suggesting the importance of coincidence detection of multiple PRRs for innate immune sensing and generating antimicrobial responses,^11^ and potentially highlighting a way to increase the strength of immunomodulatory responses. Our results confirm that synergistic interaction is also required for Pam2ODN efficacy against inflammation, as neither Pam2 nor ODN showed any efficacy when delivered alone. We believe this highlights the importance of evaluating combinations of immunomodulatory agents for future human studies. While Pam2 and ODN were the strongest combinations of PRR agonists for antimicrobial efficacy,^7^ a systematic screen has never been performed that assessed immunomodulatory strength, and it is possible that different combinations of PRR agonists might show even greater immunomodulatory strength.

We propose that these data support a stronger focus on the bi-directional signaling that occurs between systemic immune responses and mucosal tissues. We show here that allergic polarization of mucosal tissues can proceed independently of the systemic immune system. Thus, it is plausible that treatments focused on reducing type 2 polarization of the mucosal tissue might be equally as effective as broadly blocking systemic immune responses (e.g., glucocorticoids), while likely avoiding toxicities of systemic immunosuppression.

To clarify the precise cellular and molecular mechanism by which Pam2ODN modulates the type 2 immune response and reduces allergic responsiveness, further investigation is required. However, in this study we have shown that Pam2ODN is a generalizable immunomodulatory agent that counteracts allergic type 2 immune responses and is likely superior to previously tested single agents, due to synergistic cooperation. The public need for immunomodulatory agents that can counteract the morbidity of type 2 allergic disease is great, and clinical studies with Pam2ODN have already shown the drug to be well tolerated in humans (NCT02124278, NCT02566252), allowing Pam2ODN to be efficiently evaluated in the setting of allergic immunotherapy.

## Materials and Methods

### Mice

Animal studies are reported in compliance with the ARRIVE guidelines^20^ and with recommendations made by the *British Journal of Pharmacology*.^21^ Breeder BALB/cJ mice were obtained from the Jackson Laboratory (Sacramento, CA) and housed in specific pathogen-free conditions on a 12-hour light/dark cycle with free access to food and water. For euthanasia, mice were injected intraperitoneally with 2,2,2-tribromoethanol (250 mg/kg) and exsanguinated by transection of the abdominal aorta. All procedures were performed in accordance with the Institutional Animal Care and Use Committee of MD Anderson Cancer Center and the Texas A&M Institute for Biosciences and Technology. Chemicals were obtained from Sigma-Aldrich (St. Louis, MO) unless otherwise specified.

### Treatment with aerosolized Pam2ODN

This was performed as described.^15^ Briefly, ODN 5’ TCG TCG TCG TTC GAA CGA CGT TGA T 3’ as the sodium salt on a phosphorothioate backbone (ODN M362) was purchased from TriLink BioTechnologies (San Diego, CA) and 2,3-bis (palmitoyloxy)-2-propyl-Cys-Ser-Lys-Lys-Lys-Lys-OH (Pam2CSK4) as the trifluoroacetic acid salt was purchased from Peptides International (Louisville, KY). A solution of ODN (1 μM) and Pam2CSK4 (4 μM) in endotoxin-free sterile water (8 ml) was placed in an Aerotech II nebulizer (Biodex Medical Systems, Shirley, NY) driven by 10 L/min of 5% CO2 in air to promote deep breathing. The nebulizer was connected by polyethylene tubing (30 cm × 22 mm) to a 10-L polyethylene chamber vented to a biosafety hood. Mice were exposed to the aerosol for 20 min, resulting in the nebulization of ∼4 ml of O/P solution.

### Sensitization and airway challenge with HDM

Lyophilized HDM extract (Stallergenes Greer, Lenoir, NC) was resuspended in PBS. Mice were sensitized to 100 μg of HDM (protein weight) by depositing 40 μl of reconstituted HDM into the oropharynx and allowing aspiration into the lungs, with mice suspended by the upper incisors on a board at 60 degrees from horizontal under isoflurane anesthesia. Mice were then challenged with 10 µg of HDM by depositing the same volume of reconstituted HDM into the nasal vestibule. This model and HDM dosages were selected from previously published data and confirmed by us during pilot studies (Fig. S1). In one case, mice were sensitized to 100 µg of HDM by intraperitoneal injection of 40 ul of reconstituted HDM. For all experiments, mouse age at the time of sensitization to HDM was between 6 and 7 weeks old.

### Sensitization and airway challenge with OVA

Mice were sensitized to 20 g ovalbumin (OVA) (Grade V, 2.25 mg alum in saline, pH 7.4; Sigma, St. Louis, MO) by intraperitoneal (i.p.) injection on day 0 and 7. Sensitized mice were exposed for 30 min to an aerosol of 2.5% (wt/vol) ovalbumin in PBS, using an Aerotech II nebulizer, as described above.

### Airway challenge with Aspergillus oryzae and OVA

Aspergillus oryzae protease (Sigma-Aldrich, St. Louis, MO) was resuspended in PBS. Mice were challenged with 10 ug A.oryzae and 2.5% (wt/vol) OVA by aerosol 2.5 times per week for 3 weeks. A. oryzae dose was selected from previously published data.^22^ Aerosol challenge was performed as described above for OVA.

### Bronchoalveolar lavage (BAL) and differential leukocyte analysis

This was performed by instilling and collecting two 1 ml aliquots of ice-cold PBS through a 20 gauge cannula inserted through rings of the exposed trachea of euthanized animals, then combining the aliquots as described^15^. Total leukocyte count was determined using a hemocytometer, and differential counts by cytocentrifugation of 100 μl of lavage fluid at 300 g for 5 min followed by Wright-Giemsa staining.

### Histochemistry

Lungs were fixed by intratracheal inflation with 10% formalin to 20 cm H2O pressure for 12 h, and then embedded in paraffin. Tissue blocks were cut into 5-µm sections, mounted on frosted glass slides (Medline, Northfield, IL), deparaffinized with xylene, washed with ethanol, then rehydrated and stained with hemoxtylin & eosin (H&E) (Sigma-Aldrich, St. Louis, MO).

### Epithelial mucin content

Epithelial mucin content was measured as described.^14,23,24^ Lungs were fixed and processed, as described above, but stained with periodic acid fluorescent Schiff reagent (PAFS). Left-axial bronchus sections were obtained using a custom precision cutting instrument (ASI Instruments, Warren, MI), as previously described.^12^ Images were acquired by investigators blinded to mouse treatment, and morphometric analysis of the images for quantitation of intracellular mucin was performed using MATLAB Software (Mathworks Software, Natick, MD). Data are presented as the area of intracellular mucus, normalized to the length of the basement membrane.

### Statistical analysis

All data sets were first analyzed with the Shapiro-Wilk test to determine normality. For analyses where there was only a single comparison between two groups, data was analyzed by Student’s t-test or Mann-Whitney U test for normally and non-normally distributed data, respectively. For analyses where multiple experimental groups were compared against a single control, data were first analyzed by one-way ANOVA or ANOVA on ranks to determine if a significant difference between any groups was present for normally and non-normally distributed data, respectively. If a significant difference was found by ANOVA, data were further analyzed using Dunnett’s test. For analyses where each experimental group was compared to every other group, data were further analyzed using Tukey’s test for one-way ANOVA or Dunn’s test for all pairwise combinations for ANOVA on ranks. Significance was determined from adjusted P values. For studies comparing Pam2 and ODN individually and together, an interaction score from linear regression analysis was used to determine whether there was a synergistic effect. All data were analyzed using Prism (version 9, GraphPad Software, San Diego, CA). P < 0.05 was considered statistically significant.

### Nomenclature of targets and ligands

Key protein targets and ligands in this article are hyperlinked to corresponding entries in http://www.guidetopharmacology.org, the common portal for data from the IUPHAR/BPS Guide to PHARMACOLOGY^25^ and are permanently archived in the Concise Guide to PHARMACOLOGY 2019/20^26^.

## Abbreviations

AHR: airway hyperresponsiveness
TLRs: Toll-like receptors
Pam2: Pam2CSK4
ODN: ODNm362
HDM: House Dust Mite
OVA: ovalbumin
Ao: Aspergillus oryzae
PRR: pattern recognition receptors
PAMPS: pattern-associated molecular patterns
ILC2: type 2 innate lymphoid cell
DC: dendritic cell
SeV: Sendai virus
TSLP: thymic stromal lymphopoietin

## Supplementary Materials

### Funding

R35 HL144805 and DP2 HL123229 to SEE. R01 HL129795 to BFD. DLG was a Howard Hughes Medical Institute Medical Research Fellow.

### Author contributions

DLG, SEE, and BFD conceived the study; DLG wrote the manuscript; DLG created the figures; DLG, GVH, SW, VVK, MKL, AMJ, RPC, AMF performed the studies.

### Competing interests

SEE, MJT and BFD are inventors on US patent 8,883,174 “Compositions for Stimulation of Mammalian Innate Immune Resistance to Pathogens”, which has been licensed by their employer, the University of Texas MD Anderson Cancer Center, to Pulmotect, Inc., which is developing Pam2ODN as a therapeutic for respiratory infections. In addition, SEE, MJT, and BFD hold equity in Pulmotect, Inc.

## Supplementary Figures

**Figure S1.**
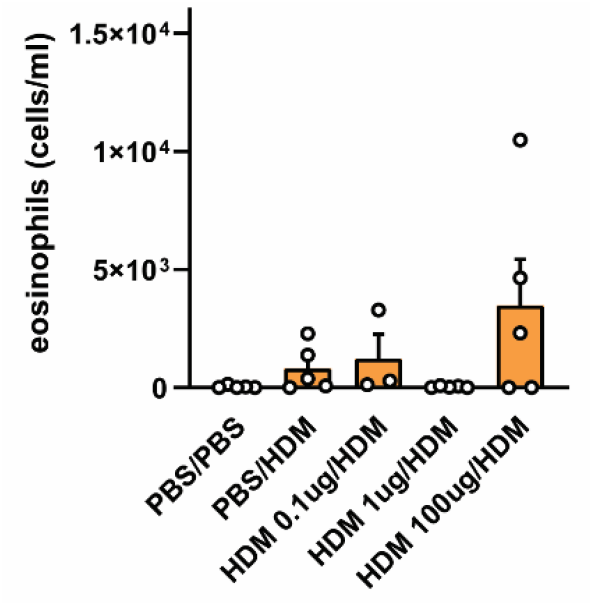
HDM pilot studies. Mice were sensitized to HDM at doses indicated, and then challenged with 10 ug HDM from day 7 to day 12. Lung inflammation was assessed at day 15.

**Figure S2.**
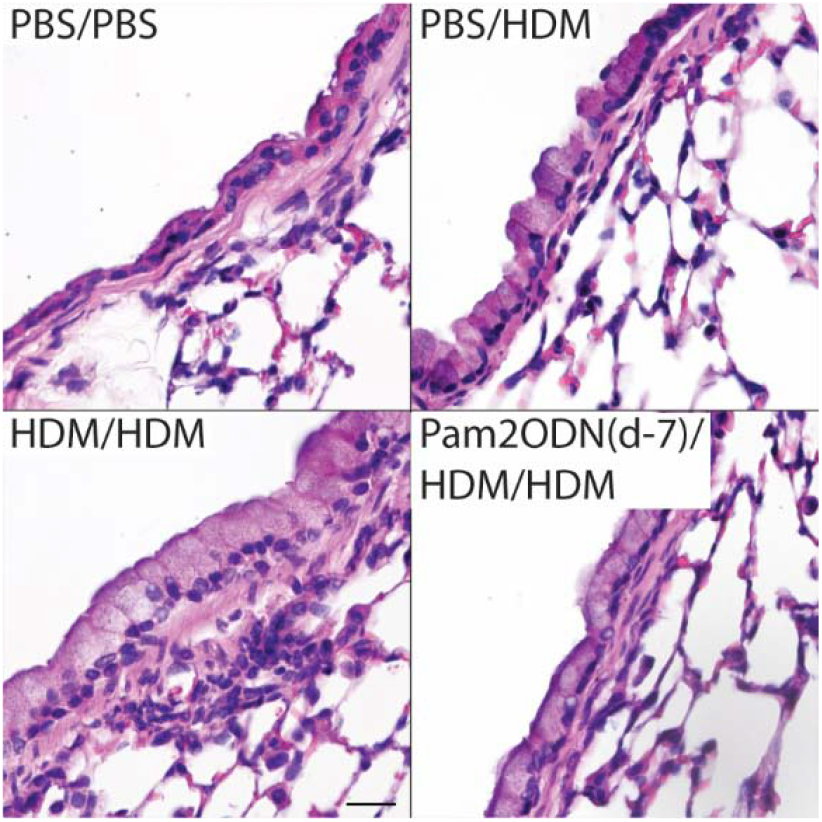
Pam2ODN attenuates allergic inflammation to HDM. Pam2ODN treatment was administered 7 days before sensitization of HDM and lungs stained for H&E to inflammation and tissue morphology. Scale bar = 20 µm.

**Figure S3.**
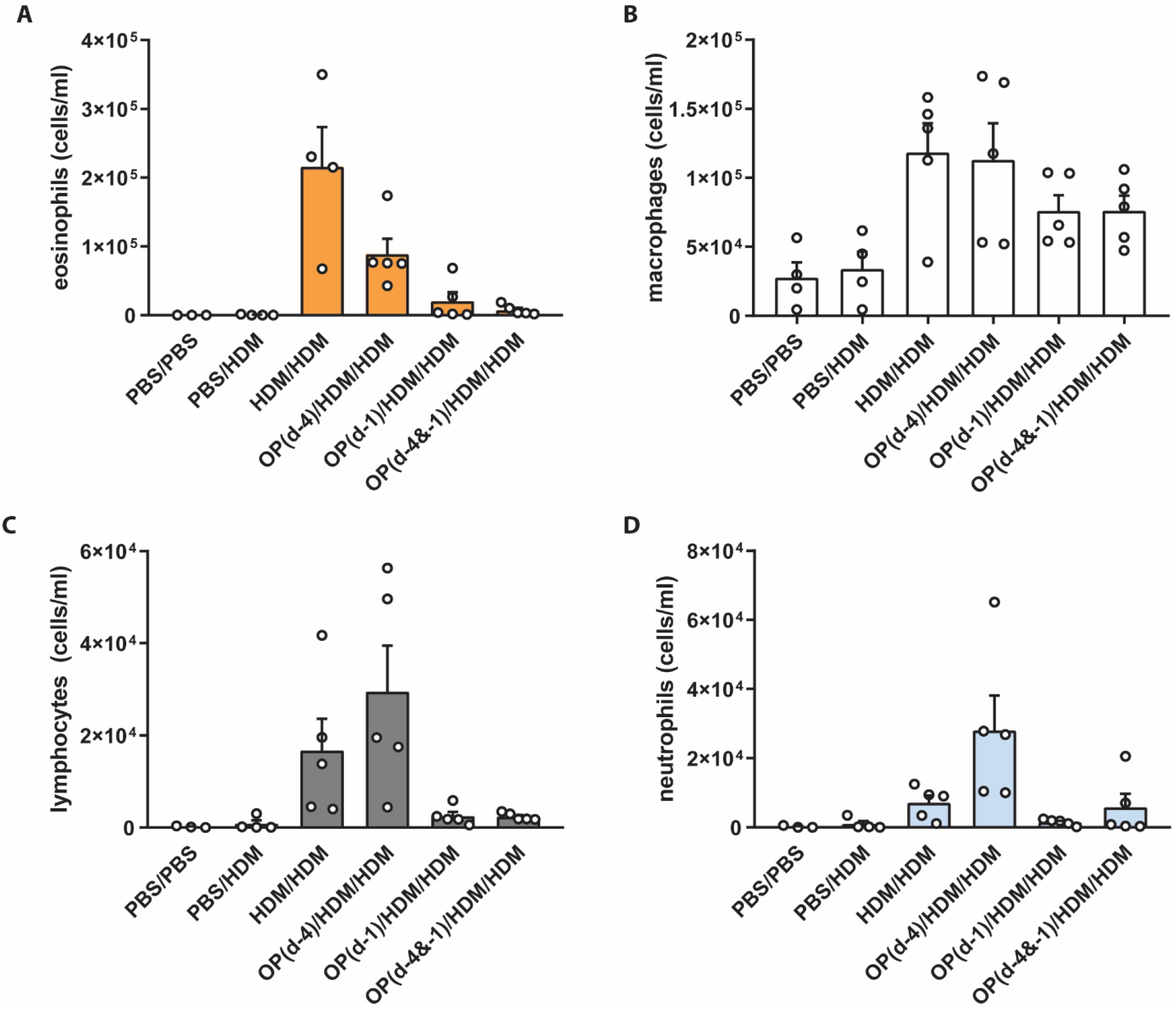
Multiple Pam2ODN treatments are better than a single treatment. Pam2ODN treatment 4 days and 1 days before HDM sensitization was superior to Pam2ODN treatment on either day alone.

**Figure S4.**
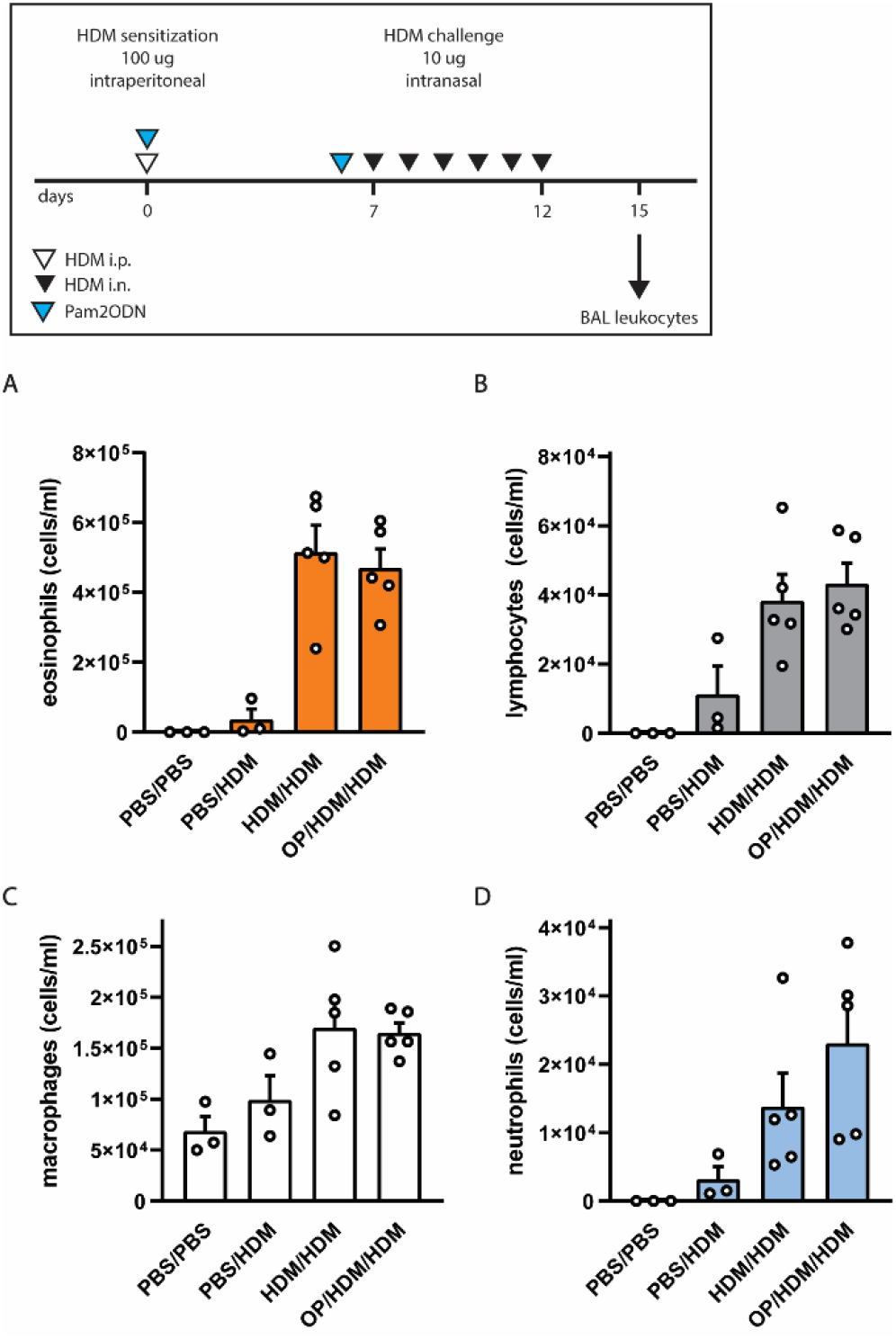
Pam2ODN does not attenuate HDM if systemically sensitized. Pam2ODN treatment on day 0 and day 7 does not prevent allergic inflammation to HDM when host is sensitized to 100 μg HDM intraperitoneally.

## References

1. Barnes, P. J. Immunology of asthma and chronic obstructive pulmonary disease. Nature Reviews Immunology 8, 183–192 (2008).

2. Lambrecht, B. N. & Hammad, H. The immunology of the allergy epidemic and the hygiene hypothesis. Nature Immunology 18, 1076–1083 (2017).

3. Stein, M. M. et al. Innate Immunity and Asthma Risk in Amish and Hutterite Farm Children. N. Engl. J. Med. 375, 411–421 (2016).

4. Schuijs, M. J. et al. Farm dust and endotoxin protect against allergy through A20 induction in lung epithelial cells. Science (80-.). 349, 1106–1110 (2015).

5. Hammad, H. & Lambrecht, B. N. The basic immunology of asthma. Cell 184, 1469–1485 (2021).

6. Evans, S. E. et al. Inhaled innate immune ligands to prevent pneumonia. Br. J. Pharmacol. 163, 195–206 (2011).

7. Duggan, J. M. et al. Synergistic Interactions of TLR2/6 and TLR9 Induce a High Level of Resistance to Lung Infection in Mice. J. Immunol. 186, 5916–5926 (2011).

8. Tuvim, M. J., Evans, S. E., Clement, C. G., Dickey, B. F. & Gilbert, B. E. Augmented Lung Inflammation Protects against Influenza A Pneumonia. PLoS One 4, e4176 (2009).

9. Clement, C. G. et al. Stimulation of Lung Innate Immunity Protects against Lethal Pneumococcal Pneumonia in Mice. Am. J. Respir. Crit. Care Med. 177, 1322–1330 (2008).

10. Evans, S. E. et al. Stimulated Innate Resistance of Lung Epithelium Protects Mice Broadly against Bacteria and Fungi. Am. J. Respir. Cell Mol. Biol. 42, 40–50 (2010).

11. Cleaver, J. O. et al. Lung epithelial cells are essential effectors of inducible resistance to pneumonia. Mucosal Immunol. 7, 78–88 (2014).

12. Goldblatt, D. L. et al. Inducible epithelial resistance against acute Sendai virus infection prevents chronic asthmaLJlike lung disease in mice. Br. J. Pharmacol. 177, 2256–2273 (2020).

13. Willart, M. A. M. et al. Interleukin-1α controls allergic sensitization to inhaled house dust mite via the epithelial release of GM-CSF and IL-33. J. Exp. Med. 209, 1505–1517 (2012).

14. Piccotti, L., Dickey, B. F. & Evans, C. M. Assessment of Intracellular Mucin Content In Vivo. in 279–295 (Humana Press, 2012). doi:10.1007/978-1-61779-513-8_17

15. Alfaro, V. Y. et al. Safety, tolerability, and biomarkers of the treatment of mice with aerosolized Toll-like receptor ligands. Front. Pharmacol. 5, 8 (2014).

16. Ch, P. & AJ, W. Allergic fungal airways disease (AFAD): an under-recognised asthma endotype. Mycopathologia (2021). doi:10.1007/S11046-021-00562-0

17. Kirtland, M. E., Tsitoura, D. C., Durham, S. R. & Shamji, M. H. Toll-Like Receptor Agonists as Adjuvants for Allergen Immunotherapy. Front. Immunol. 0, 2951 (2020).

18. Psallidas, I. et al. A phase 2a, double-blind, placebo-controlled randomized trial of inhaled TLR9 agonist AZD1419 in asthma. Am. J. Respir. Crit. Care Med. 203, 296–306 (2021).

19. Hammad, H. et al. House dust mite allergen induces asthma via Toll-like receptor 4 triggering of airway structural cells. Nat. Med. 2009 154 15, 410–416 (2009).

20. Kilkenny, C., Browne, W. J., Cuthill, I. C., Emerson, M. & Altman, D. G. Improving Bioscience Research Reporting: The ARRIVE Guidelines for Reporting Animal Research. PLOS Biol. 8, e1000412 (2010).

21. JC, M. & E, L. Implementing guidelines on reporting research using animals (ARRIVE etc.): new requirements for publication in BJP. Br. J. Pharmacol. 172, 3189–3193 (2015).

22. J, M. K. et al. A Fungal Protease Model to Interrogate Allergic Lung Immunity. Methods Mol. Biol. 1799, 1–9 (2018).

23. Tuvim, M. J. et al. Synaptotagmin 2 couples mucin granule exocytosis to Ca2+ signaling from endoplasmic reticulum. J. Biol. Chem. 284, 9781–7 (2009).

24. Evans, C. M. et al. Mucin Is Produced by Clara Cells in the Proximal Airways of Antigen-Challenged Mice. Am. J. Respir. Cell Mol. Biol. 31, 382–394 (2004).

25. JF, A. et al. The IUPHAR/BPS Guide to PHARMACOLOGY in 2020: extending immunopharmacology content and introducing the IUPHAR/MMV Guide to MALARIA PHARMACOLOGY. Nucleic Acids Res. 48, D1006–D1021 (2020).

26. Sph, A. et al. THE CONCISE GUIDE TO PHARMACOLOGY 2019/20: Introduction and Other Protein Targets. Br. J. Pharmacol. 176 Suppl 1, S1–S20 (2019).

